# Stress Combinations and their Interactions in Plants Database (SCIPDb): A one-stop resource for understanding combined stress responses in plants

**DOI:** 10.1101/2022.12.05.519235

**Authors:** Piyush Priya, Mahesh Patil, Prachi Pandey, Anupriya Singh, Vishnu Sudha Babu, Muthappa Senthil-Kumar

## Abstract

We have developed **S**tress **C**ombinations and their **I**nteractions in **P**lants **D**ata**b**ase (SCIPDb; http://www.nipgr.ac.in/scipdb.php), a compendium and interactive platform offering information on both morpho-physio-biochemical (phenome) and molecular (transcriptome) responses of plants to different combinations of stresses. To delineate the effects of various stress combinations/categories on yield in major agricultural crops, global phenome data from 939 studies was analyzed and results showed that yield was affected to the greatest extent under the abiotic–abiotic stress category, followed by the biotic–biotic and abiotic–biotic stress categories. In the abiotic–abiotic stress category, drought–heat, heat–salinity, and ozone–UV are the major stress combinations causing high yield loss in barley, wheat, soybean, and quinoa crops. In the abiotic–biotic stress category, the salinity–weed stress combination causes highest yield loss in rice crop. In the biotic–biotic stress category, the nematode–fungus combination was most detrimental, causing considerable yield losses in potato, groundnut, and sugar beet crops. Transcriptome datasets from 36 studies hosted in SCIPDb identified novel genes. Thus far, these genes have not been known to play a role in combined stress. Integretome analysis under combined drought–heat stress pinpointed carbohydrate, amino acid, and energy metabolism pathways as the crucial metabolic, proteomic, and transcriptional components in plant tolerance to combined stress. These examples illustrate the application of SCIPDb in identifying novel genes and pathways involved in combined stress tolerance. Further, we showed the application of this database in identifying novel candidate genes and pathways for combined drought and pathogen stress tolerance in Arabidopsis and chickpea. To our knowledge, SCIPDb is the only publicly available platform that provides extensive information and paves the way for advancing mechanistic understanding of plant responses to combined stresses.

## INTRODUCTION

Abiotic and biotic stresses are the major deterrents to the achievement of global food security, necessitating the urgency to develop better-adapted crops (IPCC, 2022; Mittler and Blumwald, 2010). Plants are often exposed to combinations of stresses during their life cycle, and increasing evidence highlights that stress combinations are more potent and realistic threats to plant growth and productivity than individual stresses (Atkinson and Urwin, 2012; Ahuja et al., 2010; Desaint et al., 2021; Sinha et al., 2021, Zandalinas et al., 2021a, b). Considerable information on plant stress has accumulated over the years, but our understanding of the physiological and molecular responses of plants to combined stress remains poor (Pandey et al., 2017; Mahanligam et al., 2021; Zandalinas et al., 2020a; Zandalinas and Mittler, 2022). Combined stress studies, although under-represented compared to individual stress studies, entail voluminous and highly complex information on plant response to combined stresses (Cohen et al., 2021; Zandalinas et al., 2020b; Zandalinas et al., 2021a).

A plant perceives combined stress as a new state of stress, and adaptation strategies to stress combinations are based on the interaction between the physiological and molecular responses simultaneously triggered by each stress entity independently (Gupta et al., 2016; Lopez-Delacalle et al., 2021; Pandey et al., 2017). The outcome of such interactions may be “positive” or “negative,” wherein combined stress causes less or more damage, respectively, than the individual stresses. The outcome also depends on many factors like plant age, genotype, stress intensity, duration of the stresses, and order of stress perceived by the plant, which makes combined stress more complex to understand (Mittler, 2006; Pandey et al., 2015; Zandalinas et al., 2021b). In addition, the time of imitation of the second stress since the first stress also decides the outcome of the interaction between the stresses (Choudhary et al., 2022). These responses are mediated by switching on specific pathways and processes that are unique, specific, and sometimes even contrasting from the individual stress responses. Plants also exhibit shared responses common among individual and combined stresses (Suzuki et al., 2014; Zhang and Sonnewald, 2017).

Thus, to better comprehend the complexities of combined stress responses and fill existing gaps, there is a pressing need for a pertinent database. There is no database or platform solely dedicated to combined stress. TOMRES (https://www.tomres.eu/) and Stress Combination: A New Field in Molecular Stress Research by the University of North Texas (http://biology.unt.edu/stresscombination/) are two combined stress web resources available for specific plants and for one type of stress combination, apart from individual stress databases such as STIFDB2, QlicRice, and the Arabidopsis stress-responsive gene database (Borkotoky et al., 2013; Naika et al., 2013; Smita et al., 2011). But these resources are not extensive and broad. Here, we developed the **S**tress **C**ombinations and their **I**nteractions **I**n **P**lants database (SCIPDb; htttp://www.nipgr.ac.in/SCIPdb.php), a user-friendly platform providing options to browse, search, analyze, and download data for various stress combinations studied to date. SCIPDb provides researchers easy access to combined stress-related information and tools for extracting need-based, specific information.

## RESULTS AND DISCUSSION

### SCIPDb and its key features

SCIPDb is a comprehensive collection of morphological, physiological, biochemical, and transcriptomic data on combined stresses published to date, systematically analyzed and presented in an easy-to-use interactive database and web server (Figure 1A and 1B; Supplemental Figure 1). Currently, SCIPDb hosts phenome data curated from 939 studies, covering 123 stress combinations, 118 plant species, 283 pathogenic agents (including bacteria, fungi, oomycetes, nematodes, viruses, mycoplasmas, viroids, and insects), and 7 weed species (Figure 1). From the analysis of the phenome data, 107 agronomic traits affected by various stress combinations were identified. Of these, 45 traits were mapped to the identified Trait Ontology (TO) terms. Twenty traits among the 45 were related to plant morphology and yield; 11 were related to plant physiology; and 15 were biochemical traits corresponding to changes in enzymes, metabolites, and hormone levels. These traits can be targeted for trait-based breeding programs to develop combined stress-tolerant crops. Further, gene-to-TO relationships (Pan et al., 2019) can be derived to decipher the genome-to-phenome relationships under combined stress. The transcriptome data hosted in SCIPDb is from 36 studies available in the public domain thus far, representing 58 stress combinations and 16 plant species (Supplemental Figure 2).

**Figure 1.**
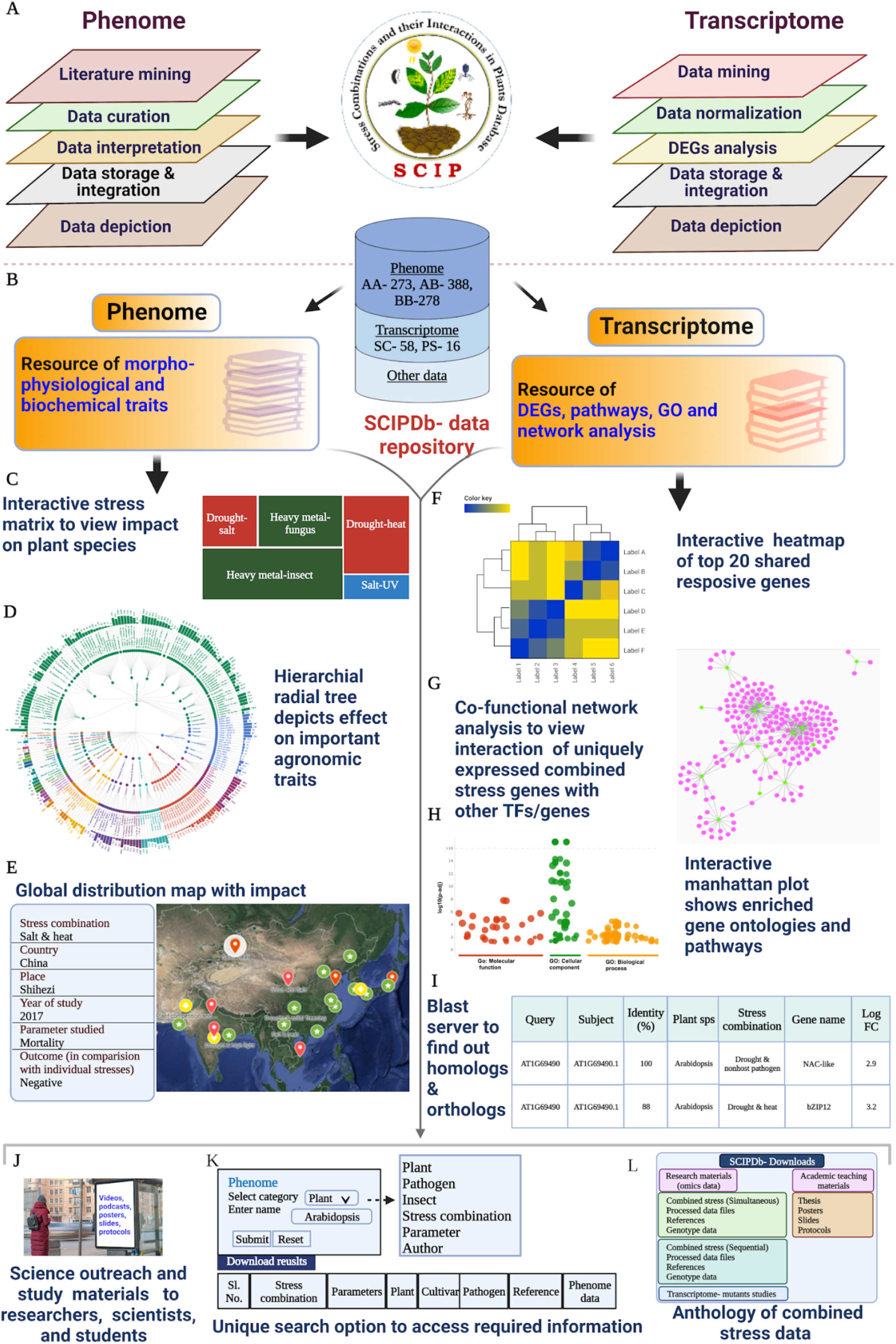
Outline of the Stress Combinations and their Interactions in Plants Database (SCIPDb), indicating its key features and applications. **(A)** The upper panel shows the steps involved in data mining, curation, analysis, and integration of phenome and transcriptome data into SCIPDb. **(B)** The lower panel shows the key features and applications offered to the users in the phenome and transcriptome sections. Orange boxes indicate the two major data sets hosted in SCIPDb. **(C)** The interactive stress matrix shows the net impact of the interaction between the stresses. The net impact of combined stress was determined by analyzing the percent reduction in plant growth, yield, and physiological traits. Three possible interactions, namely, positive (less damage under combined stress), negative (greater damage under combined stress), and others (equal damage under combined and individual stress), are depicted in green, red, and blue boxes, respectively. A stress combination is classified as positive, negative, or others based on the maximum number of studies in a particular interaction. The size of the box indicates the number of studies showing a particular interaction, i.e., a bigger size represents a greater number of studies. **(D)** The radial tree depicts the effect of individual and combined stresses on various traits in different plant species arranged in hierarchical order (starting from most to least damage). The parameters considered for developing radial trees were growth, yield, physiological, and pathogen-associated parameters. To normalize the data, percent change over control or percent change over individual stresses (in the case of pathogen-associated traits) was calculated for each trait and presented. **(E)** The interactive global map provides information on the global distribution of combined stresses and their effect on morpho-physiological, biochemical, and pathogen-associated traits. The map was generated using the geographic coordinates of the locations where the studies were conducted. **(F)** An interactive heatmap enables users to visualize the gene expression profile of the top 20 differentially expressed genes (DEGs) for a particular transcriptome. **(G)** A co-functional network depicts the correlation of the top 20 DEGs in the form of an interactive network. The co-functional network allows the user to interact with the graph, and it includes all the required gestures, including pinch-to-zoom, box selection, panning, etc., to access other metadata for each node and edge. **(H)** An interactive Manhattan plot depicts functional profiling of DEGs using various kinds of biological evidence, including Gene Ontology terms, biological pathways, and regulatory DNA elements. The X-axis represents functional terms grouped and color-coded by data sources, while the Y-axis shows the adjusted enrichment p-values in negative log10 scales. **(I)** BLAST server to find potential homologs and orthologs in SCIPDb. **(J)** SCIPDb hosts science outreach materials like posters, slides, videos, and podcasts related to combined stress, which will be useful for students, researchers, and scientists working in the area of combined stress. **(K)** The unique keyword-based search option helps to access all combined stress-related data with a single click. Searches can be performed using keywords like plant name, pathogen, insect, name of combined stress, gene ID, and gene name. **(L)** The download section provides processed phenome and transcriptome data and a reference list of combined stress articles hosted in SCIPDb. It also provides a link to mutant transcriptome studies and diverse resources related to combined stress. Numbers within the cylinder indicate the total number of articles curated and presented in the phenome and the total number of stress combinations covered under transcriptome. AA: abiotic–abiotic stress, AB: abiotic–biotic stress, BB: biotic–biotic stress, SC: stress combinations, PS: plant species. The figure was created with BioRender.com.

### Phenomics

Systematically analyzed phenomes are presented as data pages based on plant species (Supplemental Figure 3). For a holistic view of trends in the analyzed phenome data, interactive visualizations such as combined stress matrices, radial trees, and global combined stress distribution maps have been provided (Figure 1). The interactive stress matrices show the overall impact of different stress combinations on various plant species (Figure 1C). These visualizations will aid in deciphering and distilling comprehensive overviews of stress combinations across plant species, unlike from individual stresses alone. Among the 123 stress combinations, 69 combinations showed a negative impact on plant growth and productivity (Supplemental Figure 4). Twenty stress combinations showed a positive impact on plants, many of them belonging to the abiotic–biotic stress category. No combination in the abiotic–abiotic stress category showed a positive interaction, pegging abiotic stress combinations as the major threats to plant yield (Supplemental Figure 4). In 12 stress combinations, an equal number of studies reported both positive and negative impacts of combined stress on plants, with the majority belonging to the abiotic–biotic stress category (10 combinations) (Supplemental Figure 4). To decipher the impact of stress combinations on growth, yield, and physiological and pathogen-associated traits in various plant species, data were visualized in the form of a radial tree (Figure 1D). Our analysis of the different abiotic and biotic stress combinations reveals many pathogen infections that are aggravated under several concurrent abiotic stresses. It also reflects a number of pests– pathogen complexes that can pose a challenge to agricultural productivity. A global combined stress distribution map, another feature of SCIPDb, shows the prevalence of particular stress combinations in a locality with their impact on crop growth (Figure 1E). Knowledge of the occurrence of important stress combinations based on this interactive geographical distribution map can assist researchers in identifying agronomically relevant stress combinations. Our analysis of studies published from 1952 to 2021 showed a steep increase in publications about combined stress, most of which were from the Americas, Asia, and Europe, particularly from the last decade (Supplemental Figure 5A and 5B). Thus, the increase in the occurrence of combined stresses in these areas is deepening crop losses.

### Transcriptomics

Transcriptome data were analyzed and presented as interactive bootstrap tables, which enlist differentially expressed genes (DEGs) and their associated metadata in the form of KEGG pathways and genes (Supplemental Figure 6). Cross-references to important resources are provided to enable users to acquire more information directly. To further visualize the high-dimensional transcriptome data, each DEG table has been linked to interactive heatmaps, Venn diagrams, co-functional networks, and Manhattan plots (Figure 1F–1H). SCIPDb hosts co-functional networks for the top differentially expressed unique genes under multiple combined stresses. It provides speculative evidence about the genes that are co-regulated and thus might share a similar biological function or act together to control a specific phenotype. The functional annotation of DEGs acts as a key resource to elucidate the biological processes, functions, and pathways controlling various combined stresses in plants. Gene Ontology annotations provided in the form of Manhattan plots can be used to visualize enriched biological processes, molecular functions, and cellular components and pathways.

### Additional features of SCIPDb

A large fraction of genes in non-model plant species remains uncharacterized, which means that they lack functional annotation. Prioritizing candidate genes without any functional evidence in such species is challenging. The standalone BLAST server integrated with SCIPDb provides an option to query the database with batch nucleotide or protein sequences and will help users identify genes related to combined stress in the genomes of the ever-increasing repertoire of newly sequenced crop species (Figure 1I). SCIPDb hosts several videos, slides, podcasts, and protocols related to combined stress, making it a potential outreach portal to promote scientific communication and education (Figure 1J). A unique keyword-based search option allows a user to mine desired information from both phenome and transcriptome datasets (Figure 1K). SCIPDb datasets are hosted on a local FTP server, allowing users to download all curated phenomes, genotypes, transcriptomes, and references locally with just a few clicks. A user-defined download can also be done through specific sections of the database. These datasets can be further used for other downstream analyses to clearly grasp plant responses to combined stresses (Figure 1L). SCIPDb also encourages users to submit their data to the web portal to promote two-way communication and ultimately contribute to making the database a dynamic, robust, and single-stop platform for disseminating novel findings on combined stresses.

The interactive network developed by global combined stress transcriptome profiling and pathway enrichment analysis in Arabidopsis depicts the common and unique pathways between major combined stress categories hosted under the transcriptome visualization section. The “Applications” section hosts several case studies, which can help users understand how to use the diverse datasets hosted in SCIPDb to answer various biological questions about combined stress. The “References and links” section provides access to complete references of the research articles used in developing the data page, along with other related articles such as reviews, theses, and reports. A meta-phenome presents a combined trend of the net impact of stress combinations on plant performance after analyzing all the studies reported for a specific crop for a particular stress combination. Overall, these important features and tools in SCIPDb provide comprehensive information on each stress combination and can help identify the most prominent stress combination in a specific crop affecting polygenic traits like growth and yield.

### Effects of combined stress on yield and yield-attributing traits in major crops

Among the 123 reported stress combinations, 58, 41, and 24 were from the abiotic– biotic, abiotic–abiotic, and biotic–biotic stress categories, respectively (Figure 2A; Supplemental Figures 7–9). Out of the 58 stress combinations reported in the abiotic–biotic stress category, 87 studies, covering 26 plant species, were on the nematode–fungus stress combination, indicating it as one of the most evident stress combinations (Supplemental Figure 9). Global analysis of yield and yield-attributing traits belonging to plant performance, plant physiological response, and plant pathogenesis response showed greater reductions in yield under the abiotic–abiotic stress category, followed by the biotic–biotic stress category (Figure 2B). Evidently, combined drought and heat stress have caused enormous economic loss (four times greater than losses incurred by drought stress alone) amounting to ∼$200 billion in US between the year 1980-2012 (Mittler, 2006; http://www.ncdc.noaa.gov/billions/events). Several upcoming studies have indicated the role of combined drought, heat and high light in affecting plant development and metabolism (Zandalinas et al., 2020a, 2020b, 2021a). Apart from drought-heat stress combination, fungus–waterlogging, and salinity–weeds stress combinations substantially affected the yields of important monocots like wheat and barley, respectively (Supplemental Figure 10). Wheat yield in particular was greatly affected under drought–heat, drought–cold, boron deficiency–cold, and *Fusarium poae–* waterlogging stress combinations (Figure 2C). However, in the case of nematode– fungus and fungus–fungus stress combinations, the wheat yield response varied with the type of pathogen species involved in the interaction and the order of stress perceived by the plant, as shown in Figure 2C. The database also highlights several other important but lesser-known stress combinations significantly affecting plant yields. For example, in pulses and oilseeds such as peanut, cowpea, soybean, and common bean, yields were more affected under nematode–fungus, ozone–UV, fungus–insects, and drought–weeds stress combinations (Supplemental Figure 10). Among solanaceous crops, fungus (*Verticillium dahlia*) in association with nematodes (*Heterodera rostochiensis, Globodera rostochiensis*, and *Pratylenchus neglectus*) showed markedly reduction in potato yield (Supplemental Figure 10). Further, in view of understanding the aggravation of plant diseases and emergence of new disease complex, we found that biotic factors were critical in exacerbating several pathogen infections. (Supplemental Figures 11–12). In the biotic–biotic stress category, nematode (*Meloidogyne incognita* and *Heterodera indicus*) and fungus (*Fusarium udum, F. oxysporum, F. moniliforme*, and *Macrophomina phaseolina*) stress combinations were highly detrimental to maize, pigeon pea, cotton, and chickpea crops, causing more disease incidence and damage compared to individual stresses (Supplemental Figure 12). These results indicate that combined biotic stresses are more detrimental to crops than the stressors individually. In contrast, abiotic stresses have shown a positive effect in terms of reducing pathogen infection and its progression, e.g., ozone–*Phytophthora sojae* in soybean, ozone–*Bean common mosaic virus* in pinto bean, salinity–weeds in sorghum, shade–*Colletotrichum kahawae* in coffee, Mn toxicity– *Uncinula necator* in grapevine, and *Pythium myriotylum*–*R. solani* in peanut showed significant reductions in disease incidence under combined stress (Supplemental Figure 11). However, recent reviews on combined stress have also indicated that elevated drought, high temperature, and nutrient conditions make plants more vulnerable to pest or pathogen infection (Cohen and Leach, 2020; Desaint et al., 2021; Hamann et al., 2020; Savary and Willocquet, 2020).

**Figure 2.**
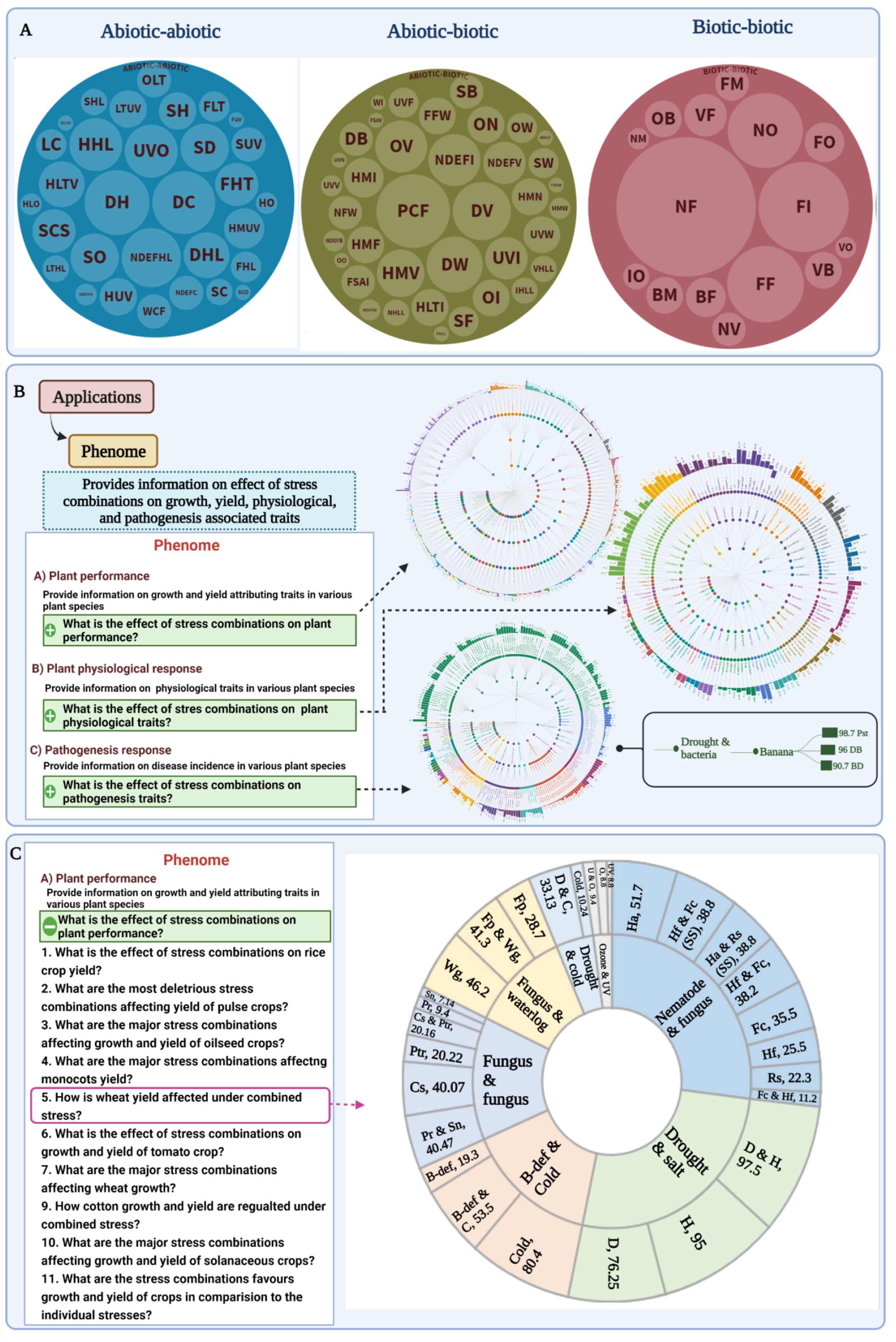
Phenome data analysis to assess the effect of stress combinations on agronomic traits. **(A)** The bubble diagrams depict the total stress combinations covered in SCIPDb under abiotic–abiotic, abiotic–biotic, and biotic–biotic stress categories. The size of the bubble is directly proportional to the number of studies under the respective stress combinations. For crop-wise stress combinations, bubble diagrams are presented in Supplemental Figures 7–9. **(B)** Schematic representation of the phenome application page and key features offered on various plant traits. Traits are classified into three major groups: plant performance (including growth and yield traits), plant physiological response (including physiological traits), and plant pathogenic response (including pathogen-associated traits). Complete information on a particular class of traits can be accessed by clicking on a particular text. The radial tree shows the overall impact of combined stress on various classes of plant traits. The tree comprises four layers: stress combination, plant species, stress treatments, and the calculated value of the trait in the form of stack bars as shown in the inset (drought and bacteria). For growth and physiological traits, values were calculated as “percent change under stress over the control,” and for pathogenesis traits, “percent change under combined stress over individual pathogen stress.” The tree should be read clockwise, where stress combinations are listed in hierarchical order based on their extent of impact on a trait, i.e., from most deleterious to least deleterious stress combination. Within a combination, stress treatments are also mentioned following similar criteria. Enlarged versions of radial trees are given in Supplementary Figures 10–12. and interactive versions are presented in database. **(C)** Representation indicating the effect of different stress combinations on wheat yield. Sunburst diagrams comprise two layers; the inner layer represents the name of the stress combination, and the outer layer represents the stress treatments with the calculated percent value. Percent change in parameter value was calculated as percent change under stress over the control. In the outer layer, the size of the box is directly proportional to the percent value, i.e., higher the percent value, bigger the box. Similarly, data for other plant species can be accessed using multiple questions listed under each group. Traits included in the plant performance group are plant height, root length, biomass, leaf number, leaf area, and yield. The figure was created with BioRender.com.

### Global combined stress transcriptome analysis in plants

Transcriptome analysis from 58 combined stress transcriptomes resulted in 45, 169 unique DEGs from 16 plant species. Functional profiling of significantly enriched DEGs revealed the involvement of genes encoding key proteins like heat-shock proteins (HSPs), Ca^2+^ signal transduction proteins, phytohormone-related genes, defense-related genes, reactive oxygen species (ROS), peroxidases, cell wall-modifying genes, and cytochrome P450 superfamily proteins. Transcription factor (TF) enrichment analysis revealed significant enrichment of dehydration response element-binding protein (DREB), ABA-responsive element-binding protein (ARF), ethylene-responsive element-binding factor (ERF), heat-shock transcription factor (HSF), NAC domain-containing protein, MYB, LOB domain-containing protein, GATA TFs, and WRKY DNA-binding protein families in the DEGs. MYBs and NAC TFs have been reported to regulate pathogen and phytohormone responses like ethylene, jasmonate, and/or salicylate (Bian et al., 2021, Vemanna et al., 2019). MYB TFs have also been reported to regulate the production of secondary metabolites during the induction of stress responses via the phenylpropanoid pathway and cell wall biosynthesis (Cao et el., 2020). They are also considered excellent candidates for broad-spectrum stress tolerance improvement in plants (Atkinson et al., 2013; Rasmussen et al., 2013, Zandalinas et al 2020a, b).

Twenty different combined stress transcriptomes were analyzed in Arabidopsis (Figure 3A), which resulted in 10,804 DEGs uniquely expressed under combined stress. Further categorization into major combined stress categories, followed by an intersection analysis revealed 3,587, 3,182, and 866 DEGs unique to abiotic–biotic, abiotic–abiotic, and biotic–biotic categories, respectively. (Figure 3B). Pathway enrichment analysis of these DEGs specific to major combined stress categories suggested several key pathway clusters consistently altered under the three major combined stress categories. This includes pathways related to the metabolism of amino acids, carbohydrates, energy, carbon, lipids, secondary metabolites, and cofactors and vitamins (Figure 3C). Pathways related to glycan biosynthesis and metabolic pathways like glycosphingolipid biosynthesis, glycosaminoglycan degradation, and N-glycan biosynthesis were unique and majorly enriched in biotic– biotic combined stress categories. Ethylene and phytochrome signaling pathways were also found to be unique to biotic–biotic combined stress categories. Genetic interactions between sugar and hormone signaling, inositol phosphate metabolism, photosynthesis, and ABC transporter pathways were found to be unique under the abiotic–biotic stress category. For the abiotic–abiotic combined stress category, glycosylphosphatidylinositol (GPI)-anchor biosynthesis, mRNA surveillance pathway, ketone body synthesis and degradation, and glucose sensing and signaling in Arabidopsis were found to be unique.

**Figure 3.**
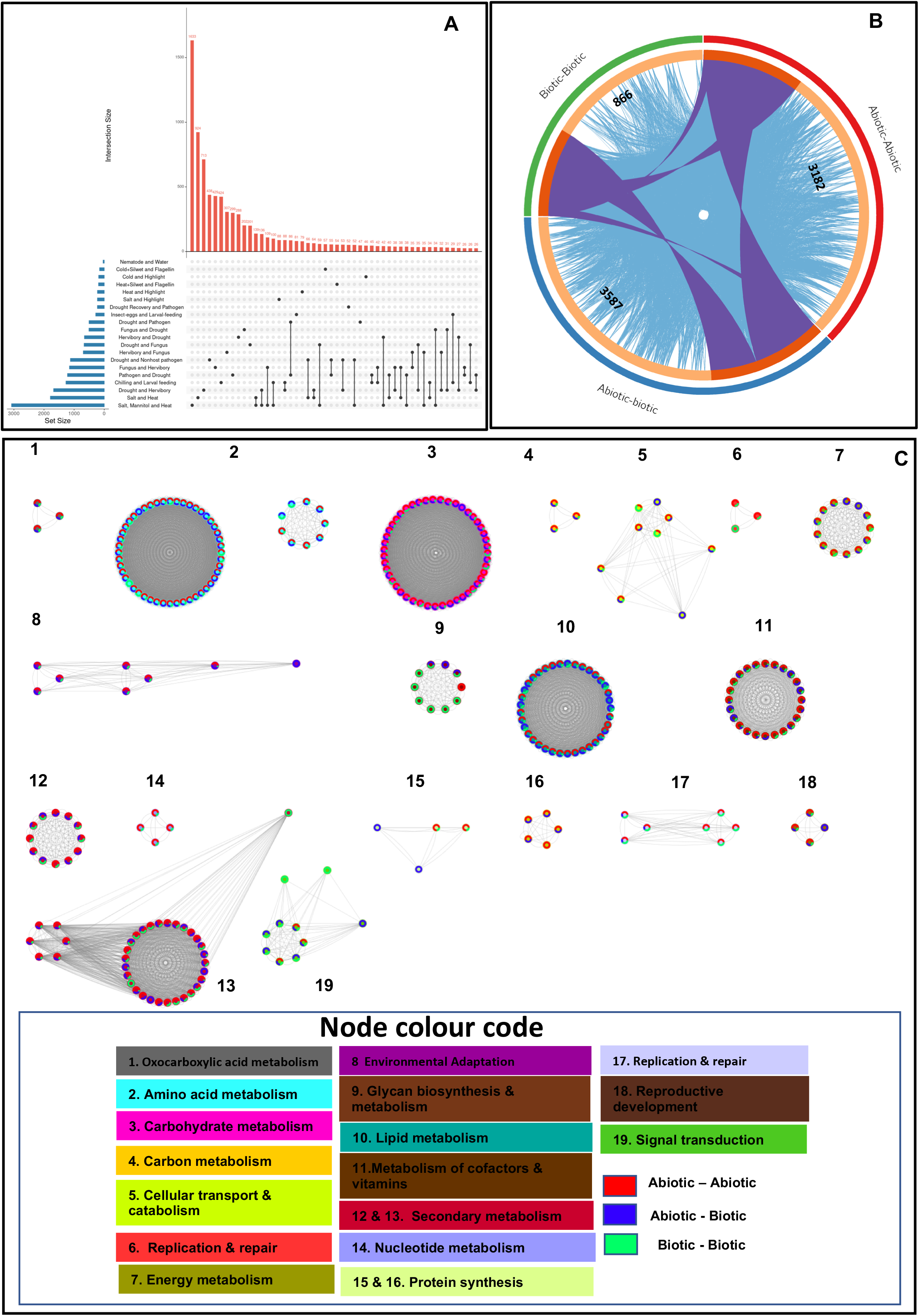
Global pathway and process enrichment analysis of differentially expressed combined stress genes in *Arabidopsis thaliana*. **(A)** Intersections among combined stress DEGs in Arabidopsis across different stress combinations. The numbers above bars indicate the number of genes within each intersection. Horizontal bars depict set size and set names. Connected dots represent common genes between the transcriptomes, while unconnected dots represent unique genes. **(B)** Circos plot representing the overlap between category-wise DEGs lists in abiotic– abiotic, abiotic–biotic, and biotic–biotic categories. The inner circle represents gene lists, where hits are arranged in the form of an arc. Genes that hit multiple lists are colored in dark orange, and genes unique to a list are shown in light orange. Purple curves link shared genes between the three categories, and blue curves link genes that belong to the same enriched ontology term. **(C)** Network representation of unique and common pathway clusters among the major combined stress categories. Analysis showed the enrichment of seven main pathway clusters, namely, lipid metabolism, carbohydrate metabolism, amino acid metabolism, biosynthesis of plant hormones, sugar and hormone signaling, secondary metabolite biosynthesis, and glycan metabolism. Ellipse-shaped nodes depicted as donuts are key pathway clusters (names indicated). The pathway clusters were grouped into broader categories based on KEGG pathway classification (for details on each node, refer to the “Transcriptome – Visualize Transcriptomics data” link in the database). Nodes in the circle represent the genes mapped to those pathways. The color of the nodes indicates the different enriched pathways and their corresponding genes in green. The network is visualized with Cytoscape (v3.8.2) with a “Group by attribute circle” layout. The network of enriched terms is represented as donut charts, where donuts are color-coded based on the identities of gene lists. The size of a donut is proportional to the total number of hits that fall into that specific term.

### Deciphering key genes and pathways under combined drought and heat stress by integrative multi-omics

While multi-omics approaches like joint pathway analysis have been limited, they are now being increasingly used in plants (Bjornson et al., 2017; Crandall et al., 2020; López-Hidalgo et al., 2018), with the underlying hypothesis that by combining evidence from multi-omics, it will be possible to concretely pinpoint the pathways involved in the underlying biological processes. Carbohydrate metabolism and related gene expression have been identified to contribute to the superior heat and drought tolerance of anthers in the rice cultivar N22 compared to the cultivar Moroberekan (Li et al., 2015). The joint pathway analysis approach, integrating changes in gene expression, proteome, and metabolite concentrations in drought and heat combined stress treatments, suggested significant enrichment of four major classes of pathways enriched based on the KEGG BRITE hierarchy. Amino acid metabolism, energy metabolism, carbohydrate metabolism, and signal transduction pathways were supported by all three omics (Figure 4A) (Zandalinas et al., 2022). Within the amino acid metabolism class, significantly enriched pathways were those related to glutathione metabolism; alanine, aspartate, and glutamate metabolism; glycine, serine, and threonine metabolism; and cysteine and methionine metabolism. Pentose phosphate pathway; glycolysis or gluconeogenesis; and pyruvate, fructose, mannose, ascorbate, aldarate, amino sugar, and nucleotide sugar metabolism pathways were found to be enriched within the energy metabolism class, while carbon fixation in photosynthetic organisms, nitrogen metabolism, and sulfur metabolism pathways were enriched under the carbohydrate metabolism class. Among the signal transduction pathways, phosphatidylinositol signaling system pathways mapped to all the three omics data analyzed. Thus, genes commonly associated between these pathway classes may play a significant role in combined stress tolerance in plants (Figure 4B).

**Figure 4.**
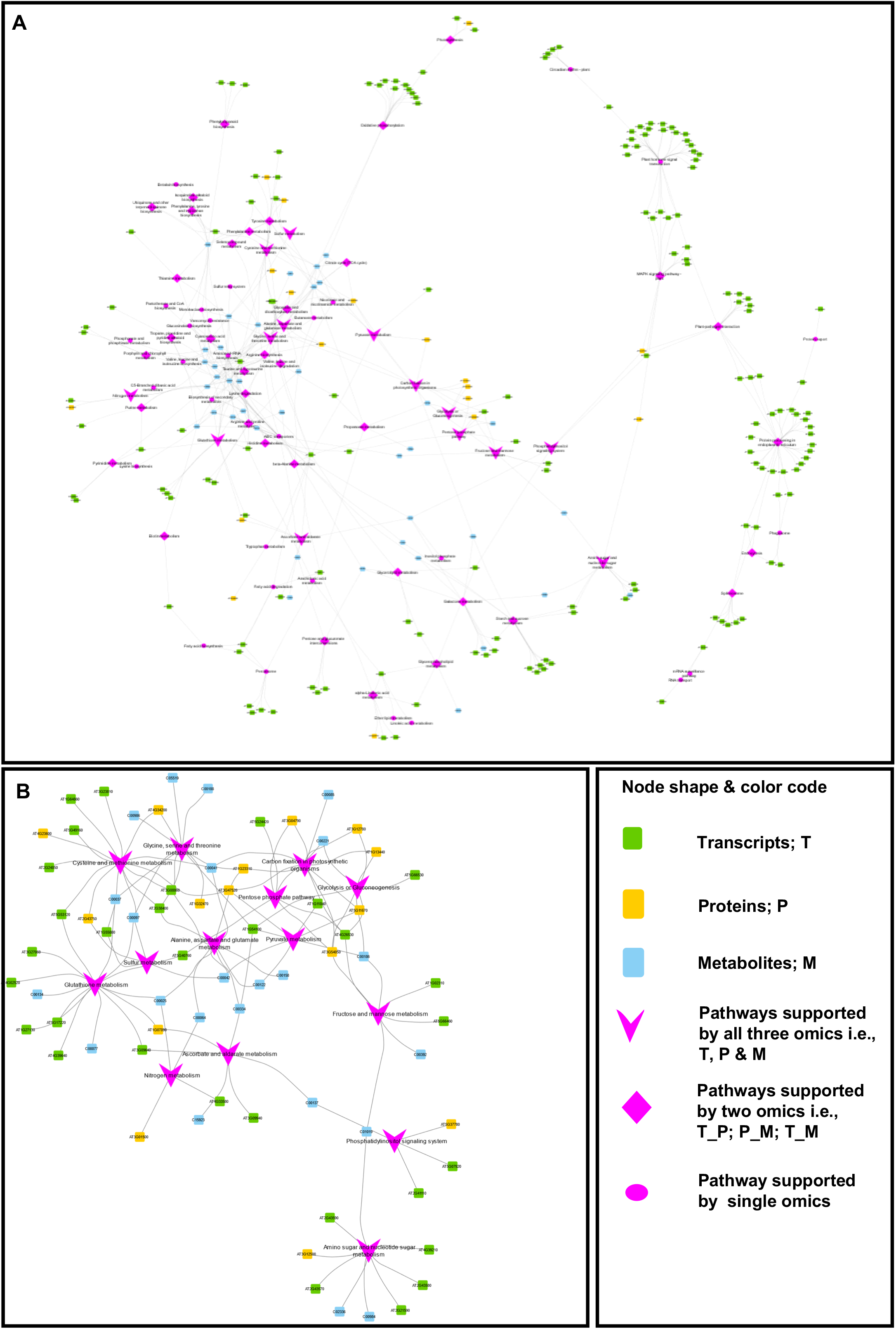
Integrative multi-omics analysis to decipher key omics features and pathways differentially altered during the drought and heat stress combination in Arabidopsis. **(A)** The network representation of differentially regulated genes, proteins, and metabolites under the drought and heat stress combination in Arabidopsis was done via joint pathway analysis and visualized in the “Edge weighted spring embedded” layout in Cytoscape (v3.8.2). The network is presented as nodes indicating various pathways and their associated omics features connected by edges. Edges have been bundled for clarity. The size of a pathway node represents the pathway impact in terms of evidence from omics features, wherein arrow-headed nodes signify pathways having evidence from all the three omics features (transcriptome (T), proteome (P), and metabolome (M)), while diamond-shaped nodes have evidence from either of the two omics features (T-P, P-M, or T-M). Nodes shown in the ellipse represent pathways that show evidence from single omics (T, P, or M). Node color corresponds to the class of the pathway or features as mentioned in the node shape and color code box. **(B)** An enlarged and detailed version of the network highlighting pathways supported by all the three omics features (T, P, and M). T: transcriptomics, P: proteomics, and M: metabolomics.

### Future perspectives

Global phenome data analysis shows that abiotic–abiotic stress combinations are major threats to crop productivity. In the face of global climate change, the occurrence of these stress combinations is projected to increase in coming years. Therefore, dedicated studies on this aspect are essential to sustain crop yields in the future. Key takeaways from yield analyses are that monocots will be more affected under the abiotic–abiotic and abiotic–biotic stress category, whereas pulses, oilseeds, and vegetable crops will be more affected under the biotic–biotic stress category.

Our unique combined stress integretome developed using multi-omics data integration highlights sugar metabolism, energy metabolism, and amino acid metabolism as the key pathways operating under combined stress conditions. The addition of proteomics and metabolomics data to the database with multi-omics analysis will further demystify combined stress responses of plants. Unraveling the mechanisms by which the molecular signatures associated with these pathways impact plant response to combined stress can open new vistas for developing resilient crop varieties with better adaption to changing climate and global warming.

SCIPDb is a comprehensive database amenable to data mining and data-driven research on combined stresses in plants. With the continual accumulation of available data in the field of combined stress, we will update the database annually by incorporating newer studies. We intend to add other omics datasets related to combined stress research in the future version of SCIPDb, together with novel features like prediction modeling based on machine learning and meteorological data integration with geographical distribution information. Overall, SCIPDb is an informative and valuable resource for combined stress research in plants.

## METHODS

### Data acquisition

SCIPDb hosts two major omics datasets: phenomics and transcriptomics. For the acquisition of both the datasets, widely used search engines and public databases were extensively mined.

### Data mining for phenomics

To retrieve all available articles and to have greater than 90% literature coverage, several search engines were queried using suitable and carefully designed keywords (including several variants). Bibliography from each article was also searched to achieve better coverage.

### Data mining for transcriptomics

The relevant transcriptome datasets for combined stress in plants were compiled and curated using two major public databanks for microarray data, including Gene Expression Omnibus (GEO) (https://www.ncbi.nlm.nih.gov/geo/) and Array Express (http://www.ebi.ac.uk/arrayexpress/). The NCBI GEO and ArrayExpress functional genomics repository were queried using a large number of keywords as listed in the database. For the compilation of RNA-seq transcriptomics data, the NCBI Sequence Read Archive (SRA) (https://www.ncbi.nlm.nih.gov/sra) database was used.

### Database implementation

The frontend user interface was implemented using HTML5, CSS, and PHP (version: 7.0.12). The back-end schema was designed using MySQL, an open-source relational database management system, and data were stored in MySQL tables (Version: 5.7.17). To provide an interactive interface and enhanced user experience, we used Bootstrap 4, JavaScript, and jQuery. SCIPDb has been deployed in an Apache web server that runs on the CentOS Linux 7 (Supplemental Figure 1).

### Arabidopsis combined stress transcriptome

The upset plot was generated using the UpSetR package, while the circos plot was generated using Metascape, a gene annotation and analysis resource. (https://metascape.org/gp/index.html#/main/step1). Pathway enrichment analysis was done using major pathway databases like KEGG (https://www.kegg.jp/kegg/rest/keggapi.html), Aracyc (https://plantcyc.org/typeofpublication/aracyc), and Wikipathways (https://www.wikipathways.org/index.php/WikiPathways). Final visualization and network analysis were done using Cytoscape, v3.8.2 (https://cytoscape.org/).

### Integretome analysis

MetaboAnalystR package was used to perform joint pathway analysis of the transcriptome, proteome, and metabolome profiles. For enrichment analysis (ORA) hypergeometric analysis was used, while for topology measure, degree centrality was used. Combining p-values at the pathway level was used for the integration of the three omics datasets.

## SUPPLEMENTAL FIGURES

**Supplemental Figure 1**. Content and construction of SCIPDb.

**Supplemental Figure 2**. Combined stress transcriptome articles were analyzed and integrated into SCIPDb.

**Supplemental Figure 3**. A typical data page entry for phenome in SCIPDb.

**Supplemental Figure 4:** The heat map depicting the various stress combinations of potential environmental stresses that affect crops in the field.

**Supplemental Figure 5**. Literature analysis of combined stress articles published from 1950 to 2021.

**Supplemental Figure 6**. A typical data page entry for transcriptome in SCIPDb and its associated visualizations.

**Supplemental Figure 7**. Literature analysis of combined stress articles published from 1950 to 2021 under the abiotic–abiotic stress category.

**Supplemental Figure 8**. Literature analysis of combined stress articles published from 1950 to 2021 under the abiotic–biotic stress category.

**Supplemental Figure 9**. Literature analysis of combined stress articles published from 1950 to 2021 under the biotic–biotic stress category.

**Supplemental Figure 10**. Literature analysis of combined stress growth and yield data on various plant species.

**Supplemental Figure 11**. Literature analysis of combined stress physiological data on various plant species.

**Supplemental Figure 12**. Literature analysis of disease incidence data under combined stress on various plant species.

## AUTHOR CONTRIBUTIONS

M.S-K. conceived the idea, designed the database, outlined the manuscript, and provided all resources. Pi.P. developed the webtool and performed data integration and data visualization. M.P. and Pr.P. contributed to the phenomics part. Pi.P., M.P., and Pr.P. performed overall data analysis. A.S. contributed to the data collection for the phenomics part of the manuscript. Pi.P. and V.S.B. contributed to the data collection and analysis part of transcriptomics. M.S-K., Pi.P., M.P., and Pr.P. drafted the manuscript. M.S-K. edited and finalized the manuscript and SCIP database. All authors agreed to the submitted version of the manuscript.

## ACKNOWLEDGMENTS

The authors are thankful to the Department of Biotechnology (DBT)-eLibrary Consortium, India, and the NIPGR library for providing access to e-resources. Computational facilities provided by Genome Analysis Facility and the DBT-DISC facility at NIPGR are also duly acknowledged for sharing resources. We acknowledge Dr. Jyoti Singh for her contribution in collecting data for the phenome part.

---

**Supplemental Figure 1.**
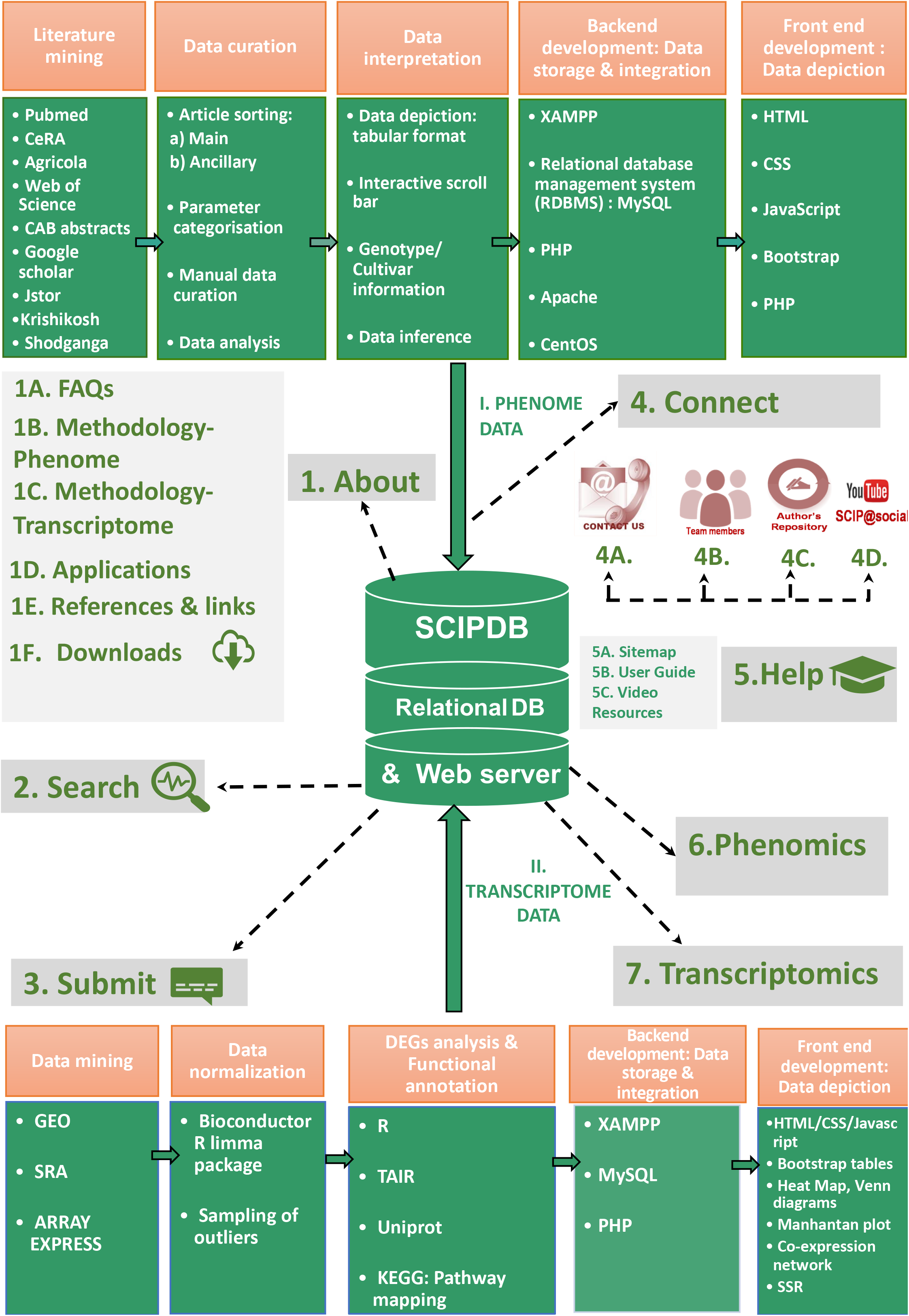
Content and construction of SCIPDb. SCIPDb provides a user-friendly interface to browse, search, download and visualize stress combination data of plants. The figure outlines the schematic representation of SCIPDb main menus and submenus. About, Search, Submit, Connect, Help, Phenomics, and Transcriptomics, are the main menus. **(1)** About menu further is subdivided into **1A. FAQs**: address common concerns, and questions, that users have. **1B. Methodology Phenome & 1C. Transcriptome:** outlines and details the steps followed to collect, curate and interpret the phenomics and transcriptomics dataset integrated into SCIPDb, **1D. Applications**: depicts the multifarious uses of SCIPDb, **1E. References & Links**: provides easy access to the entire list of research articles used in developing data pages along with other related articles such as reviews, thesis, and reports. Weblinks of labs and scientists, important books, and articles on combined stress are also provided. **1F. Downloads**: section catalogs the entire list of raw data files, references, genotypes covered in SCIPDb, along with several academic teaching materials, which can be downloaded by the user locally using the FTP server hosted hereby. (**2) The Search** menu provides the user an option to query the SCIPDb dataset based on keywords and sequence to fetch relevant information. (**3) Submit** menu provides the users with an option to submit combined stress data on phenome and transcriptome to SCIPDb. (**4) Connect** section provides the user information about **4A. Contact details, 4B. Team members, 4C. Author repository**, that provides details of authors working in the area of combined stress and **4D. SCIP@Social**, which hosts several videos and podcasts related to the area of combined stress. (**5) Help**, hosts **5A**. that further details each section and tabs of SCIPDb, **5B**. User guide: Detailed tutorial explaining steps needed to easily navigate and use SCIPDb **5C**. Video resources: Videos related to combined stress in plants. (**6) Phenomics:** Hosts morphological, physiological, and biochemical data associated with various stress combinations. (**7) Transcriptomics**: hosts a comprehensive collection of combined stress-responsive differentially expressed genes (DEGs) identified in publicly available transcriptomic data from various plant species.

**Supplemental Figure 2.**
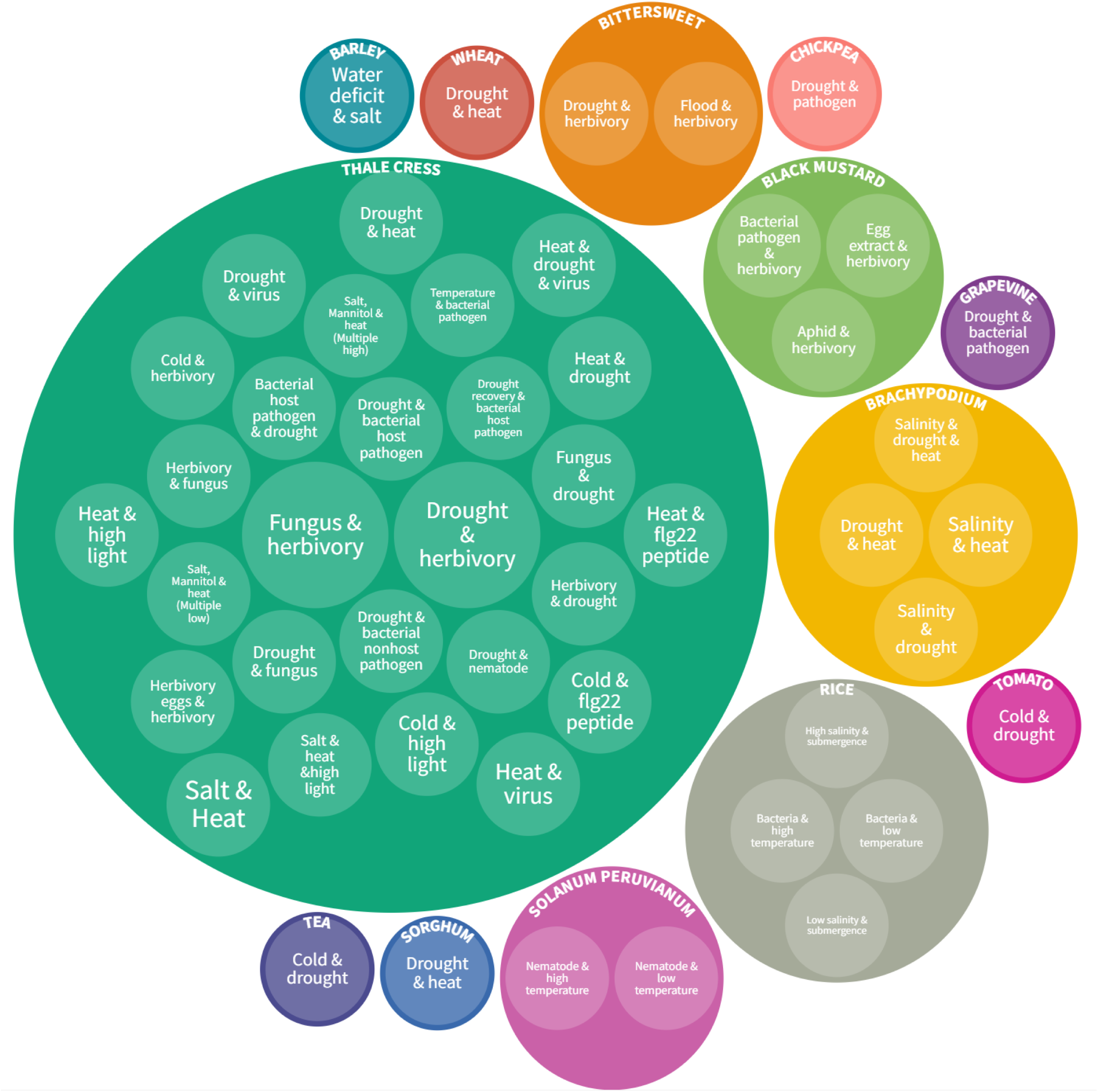
Combined stress transcriptome articles were analyzed and integrated into SCIPDb. The bubble diagram has been color-coded based on plant species and has been organized hierarchically into two layers where the first layer represents the plant species and the second layer represents the stress combinations. The size of the bubble is directly proportional to the number of articles i.e., the bigger the size more the number of studies in that stress combination for that plant species.

**Supplemental Figure 3.**
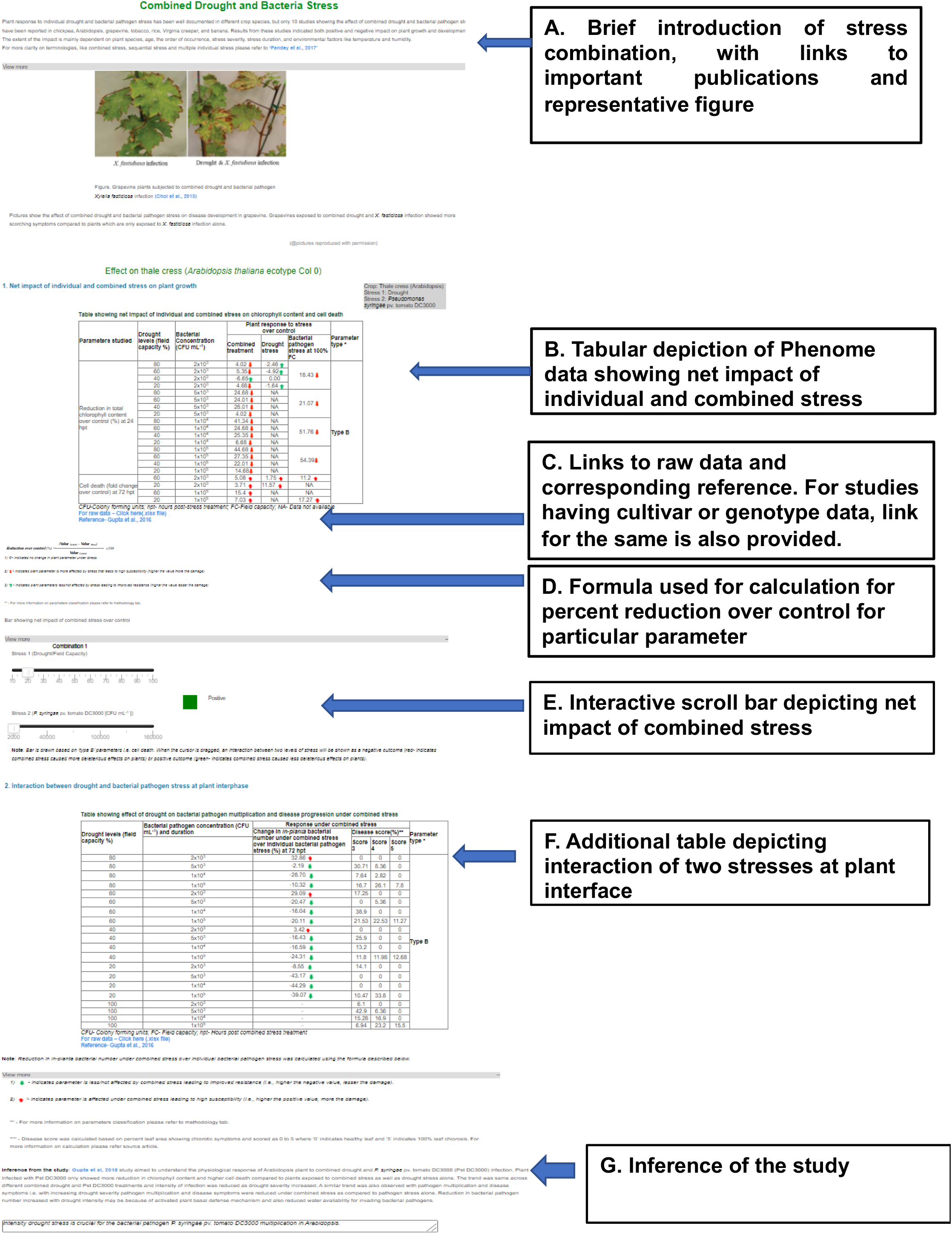
A typical data page entry for phenome in SCIPDb. The figure shows the analyzed phenome data integrated into SCIPDb. The phenome data page is organized and presented based on major stress categories, stress combinations, and plant species selection by the user. A-G details various components of the phenome data page integrated into the SCIPDb.

**Supplemental Figure 4:**
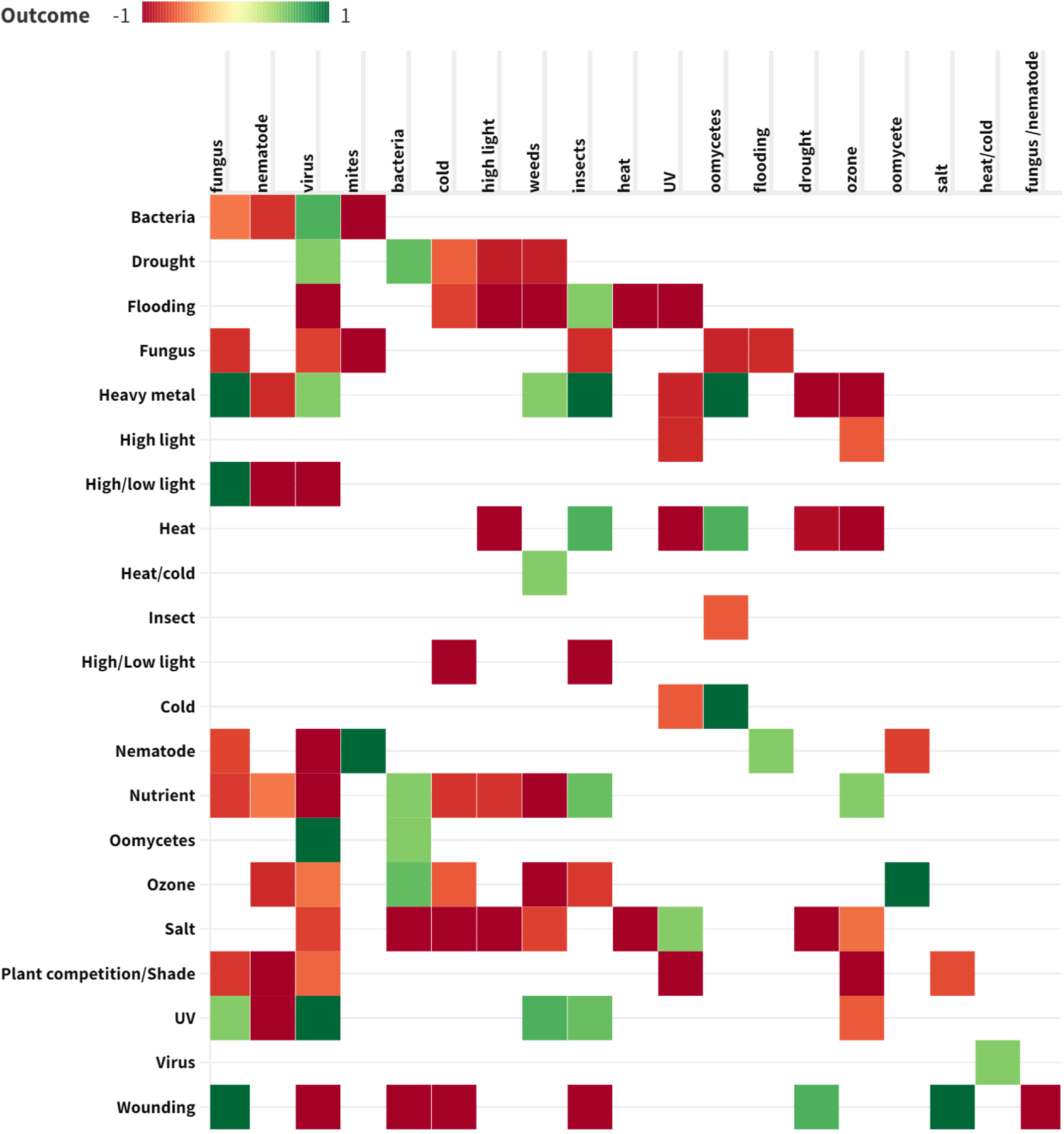
The Heat map depicting the various stress combinations of potential environmental stresses that affect crops in the field. The gradation in the color depicts potential outcomes based on findings of many studies analyzed in SCIPDb for each stress combination. Red color (−1) shows potential negative outcome, i.e, plants under these combined stresses are affected to a greater extent compared to individual stresses while green color (+1) shows potential positive outcome implying plants under these combined stresses are less/equally affected as compared to one or both the individual stresses.

**Supplemental Figure 5.**
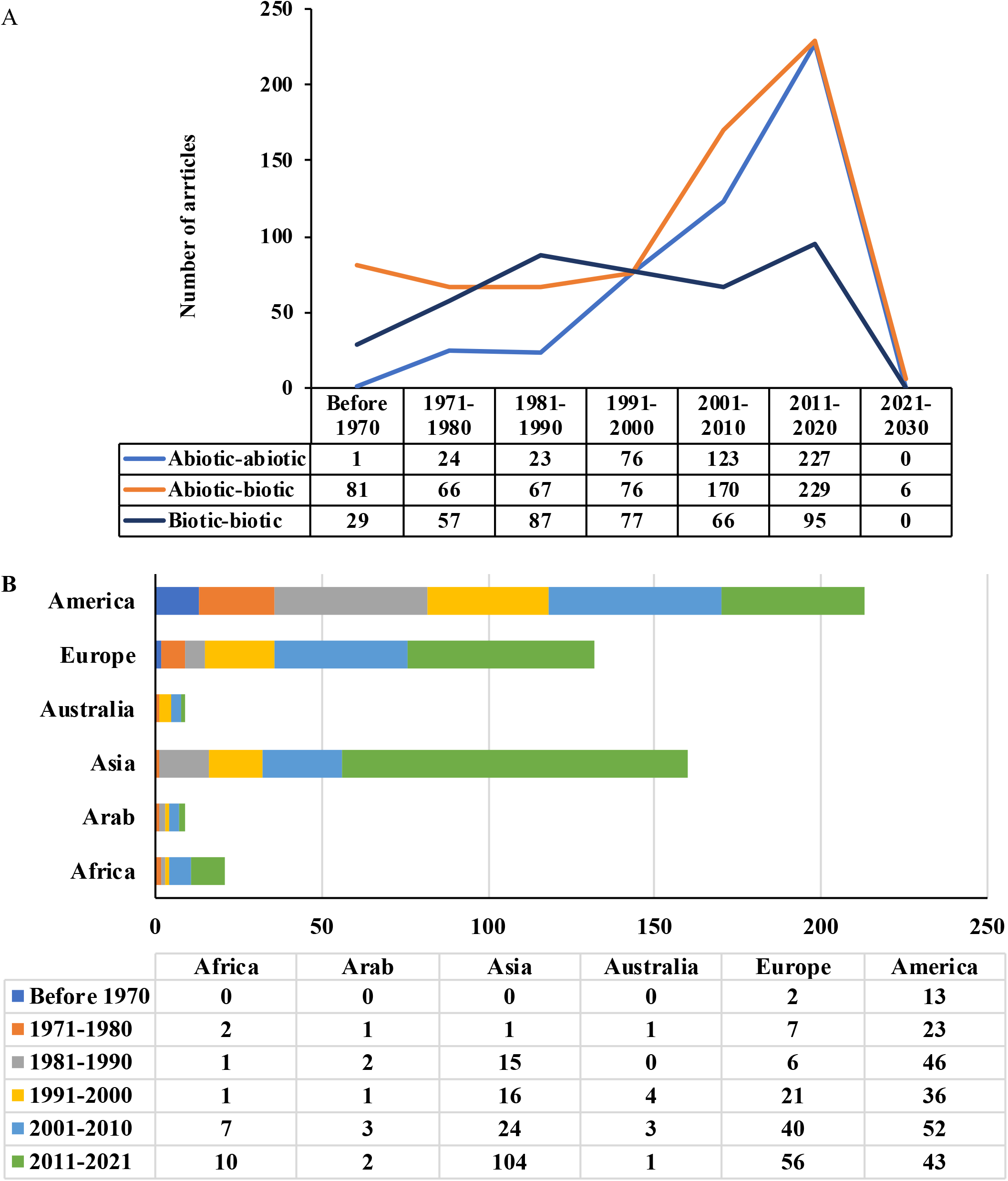
Literature analysis of combined stress articles published from 1950 to 2021. A. Graphs show the year-wise total number of articles published until the year 2021 under the abiotic-abiotic, abiotic-biotic, and biotic-biotic stress categories. B. Graph shows the year-wise distribution of combined stress articles published in eight major regions of the world. Only research articles were considered in generating these figures other types of articles like reviews, reports, mutant/transgenic studies, and articles on tree species were excluded.

**Supplemental Figure 6.**
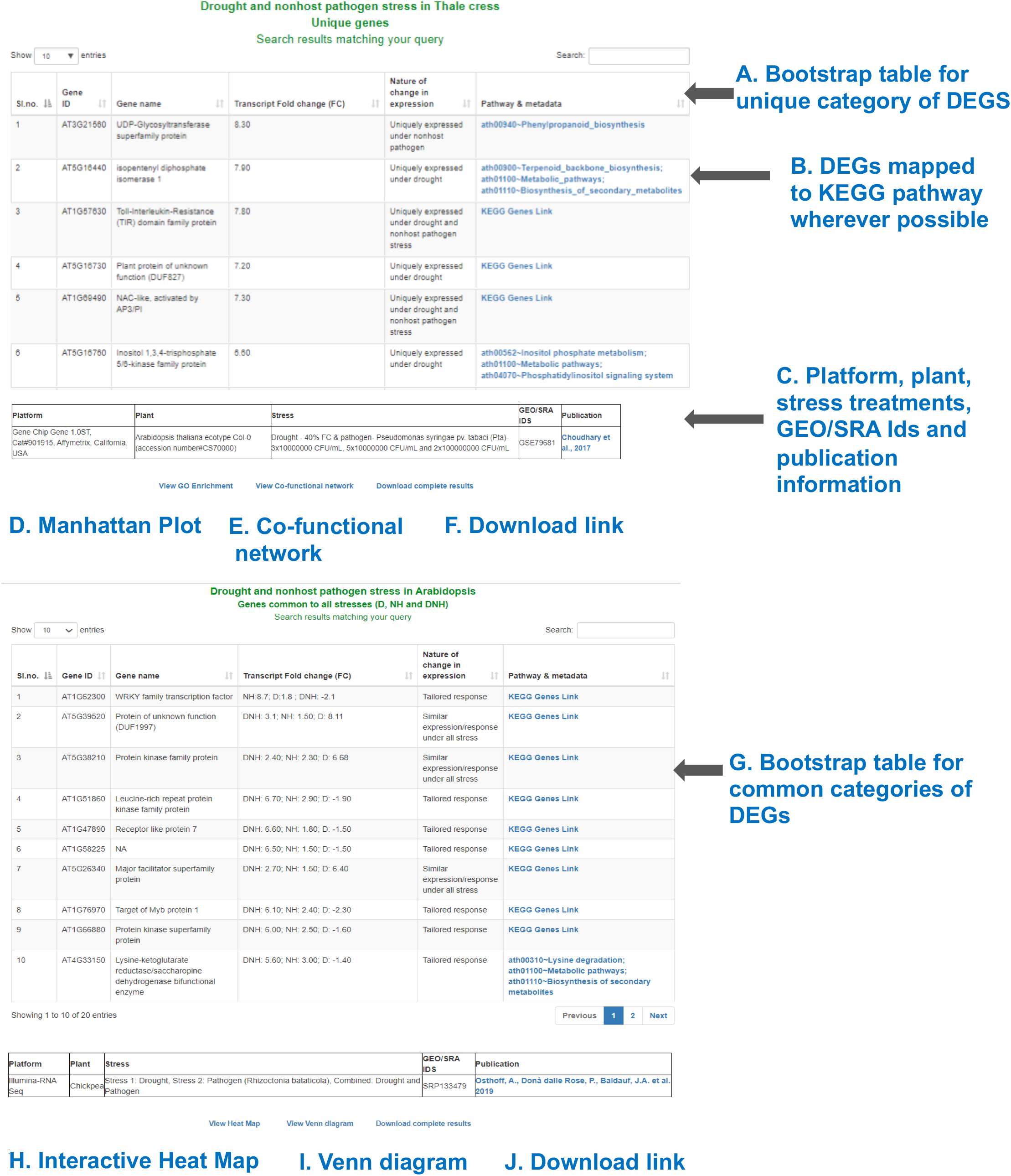
A typical data page entry for transcriptome in SCIPDb and its associated visualizations. The figure shows the analyzed transcriptome data represented in the form of an interactive bootstrap table, showing a list of DEGs, gene name, log FC, and associated metadata in the form of KEGG pathways and genes. The transcriptome data page is organized and presented based on plant, stress combination, and DEGs category selection by the user. A-J details various components of the transcriptome data page integrated into SCIPDb.

**Supplemental Figure 7.**
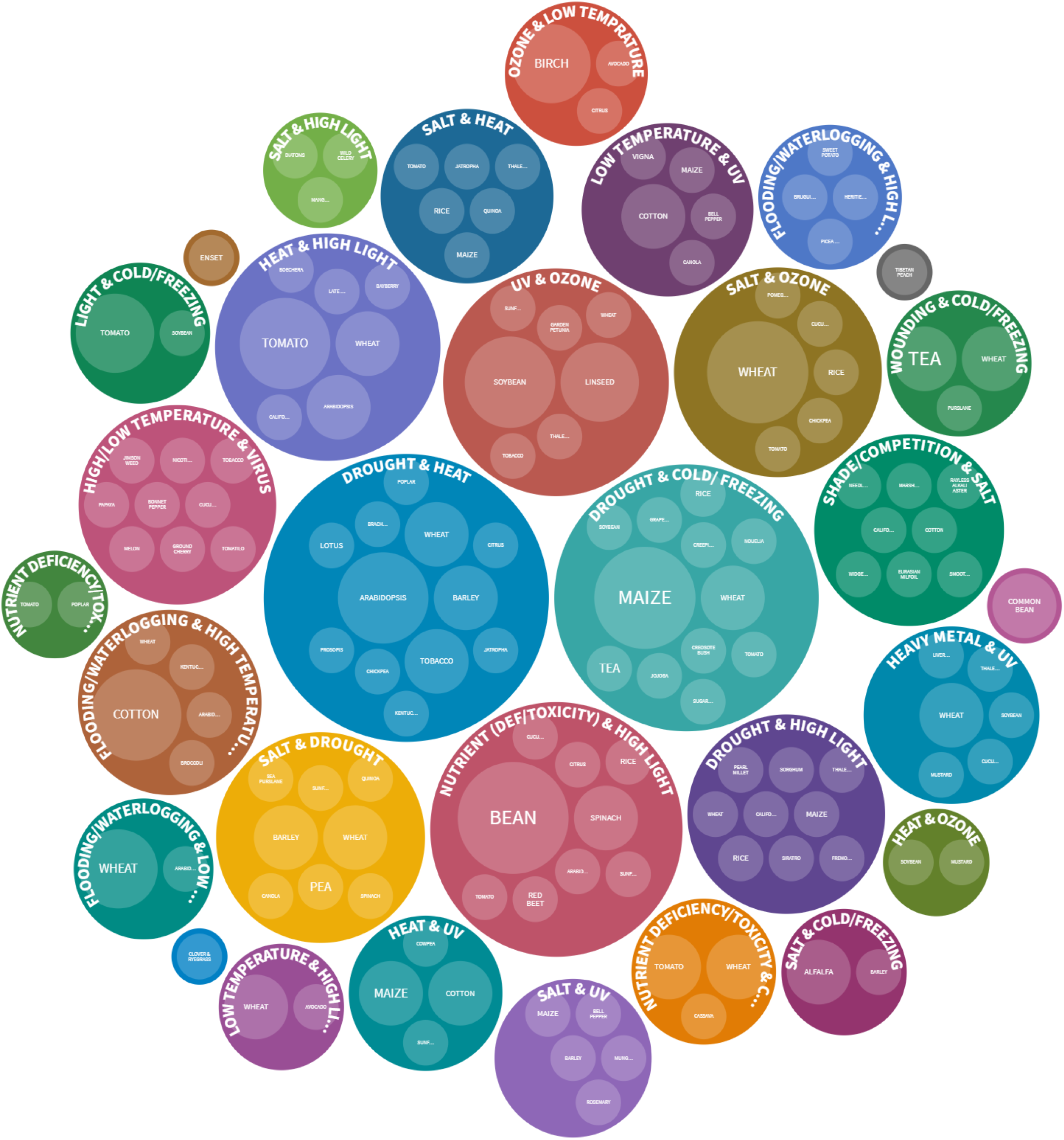
Literature analysis of combined stress articles published from 1950 to 2021 under the abiotic-abiotic stress category. The bubble diagram shows the list of stress combinations with plant species studied under the abiotic-abiotic stress category. Each bubble represents stress combinations and the size of the bubble is directly proportional to the number of articles i.e. bigger the size more the number of studies in that stress combination or plant species.

**Supplemental Figure 8.**
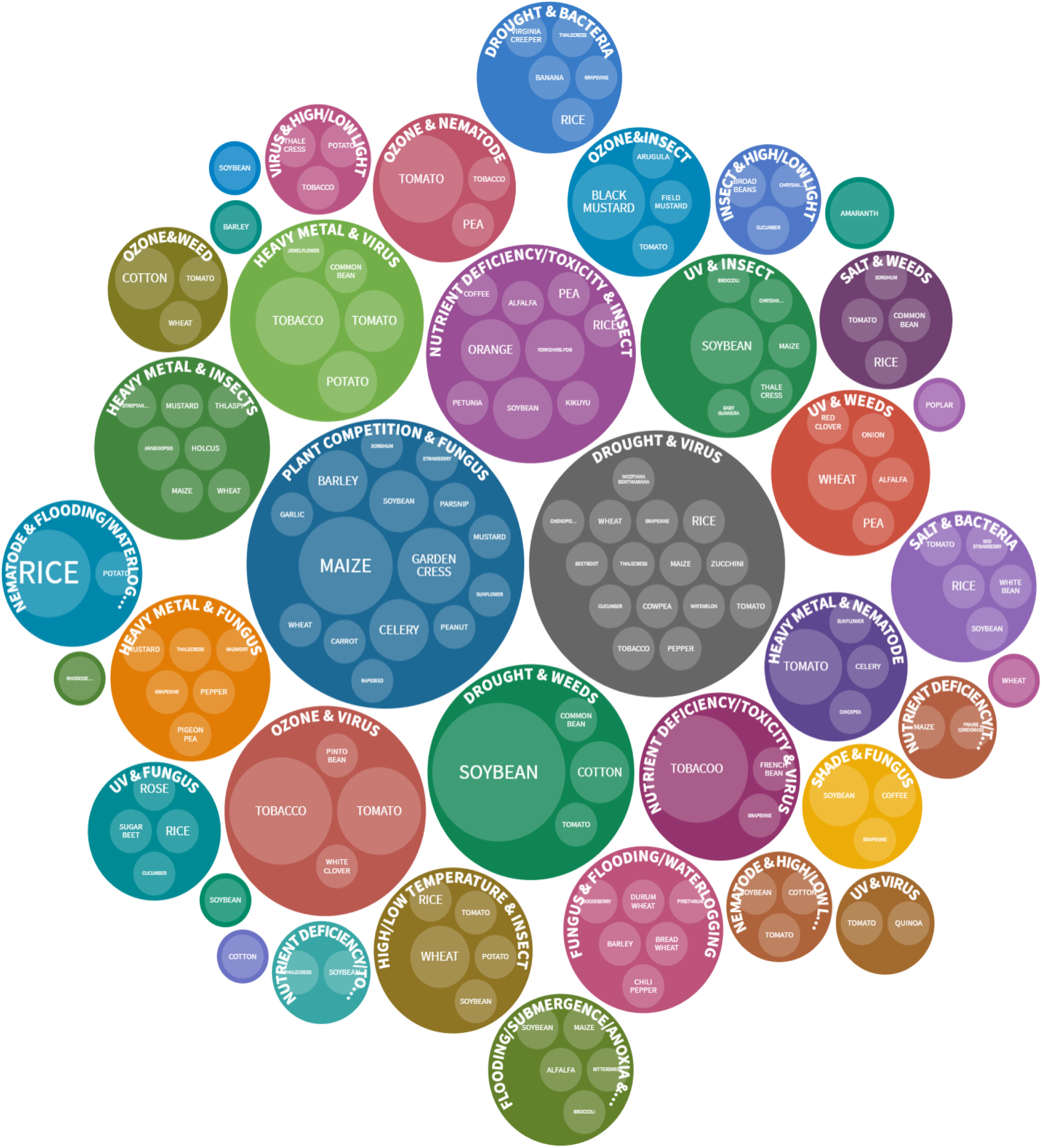
Literature analysis of combined stress articles published from 1950 to 2021 under the abiotic-biotic stress category. The bubble diagram shows the list of stress combinations with plant species studied under the abiotic-biotic stress category. Each bubble represents stress combinations and the size of the bubble is directly proportional to the number of articles i.e., the bigger the size more the number of studies in that stress combination or plant species.

**Supplemental Figure 9.**
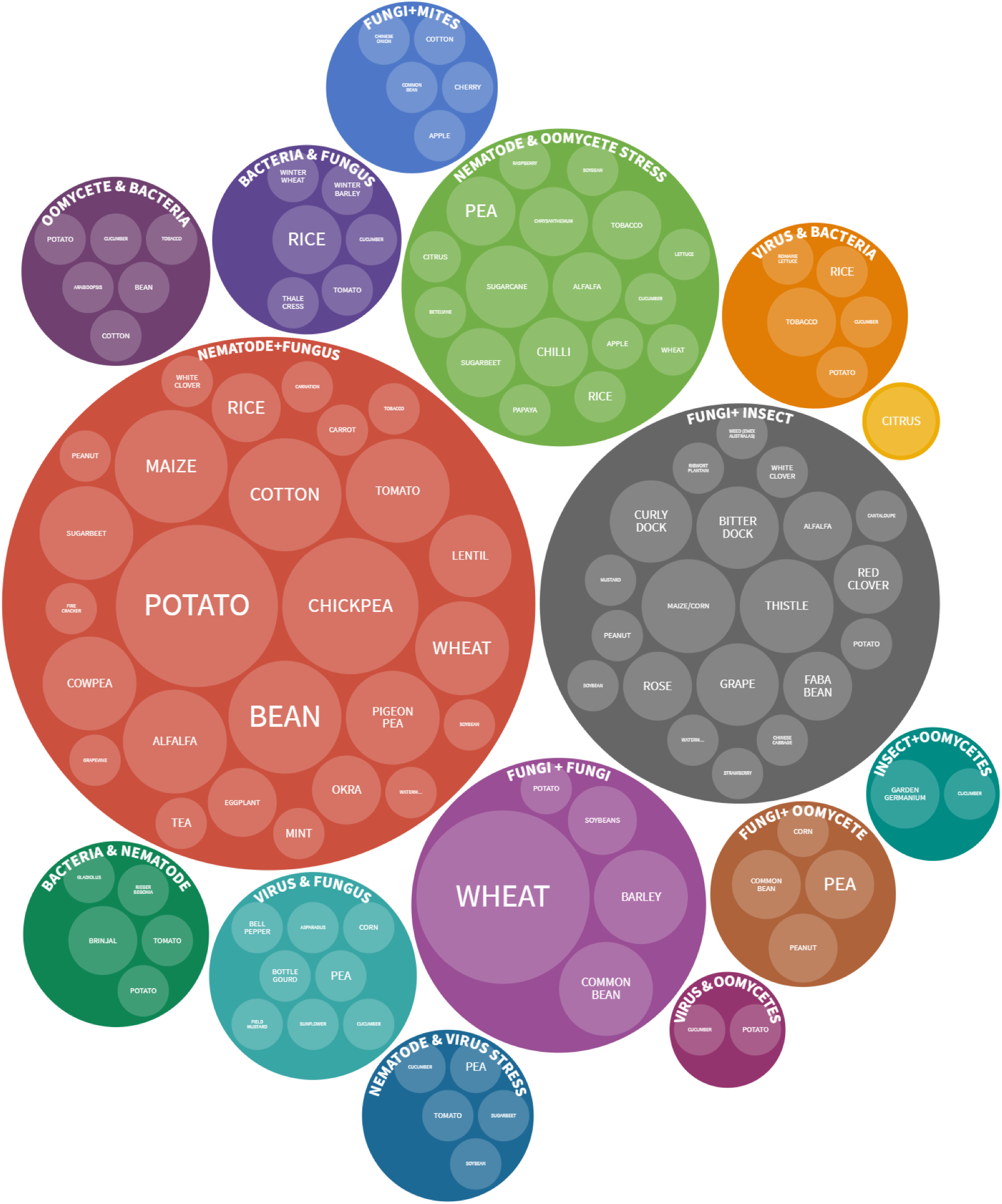
Literature analysis of combined stress articles published from 1950 to 2021 under the biotic-biotic stress category. The bubble diagram shows the list of stress combinations with plant species studied under the biotic-biotic stress category. Each bubble represents stress combinations and the size of the bubble is directly proportional to the number of articles i.e., the bigger the size more the number of studies in that stress combination or plant species.

**Supplemental Figure 10.**
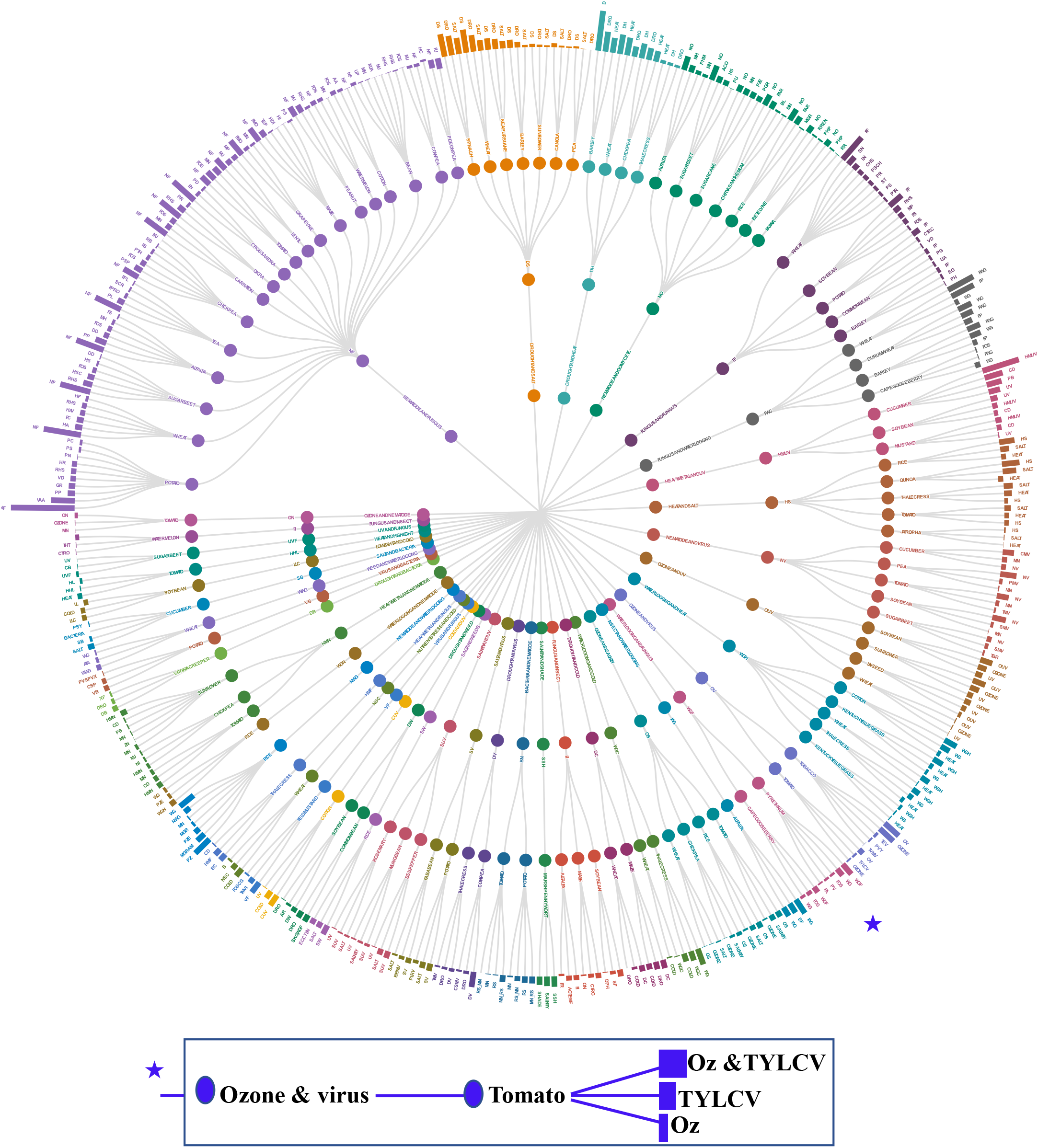
Literature analysis of combined stress growth and yield data on various plant species. The radial tree shows the effect of stress combination on growth and yield attributing traits on various plant species. The tree was developed using Flourish studio (https://flourish.studio) and Tidyverse package in R(https://www.tidyverse.org/packages/). Traits included are plant height, leaf area, leaf number, shoot weight, biomass, root weight, root length, seed weight, seed number, and yield. Percent under stress over control was calculated and using those values tree was developed. An interactive view of this tree is given on the SCIPDb website.

**Supplemental Figure 11.**
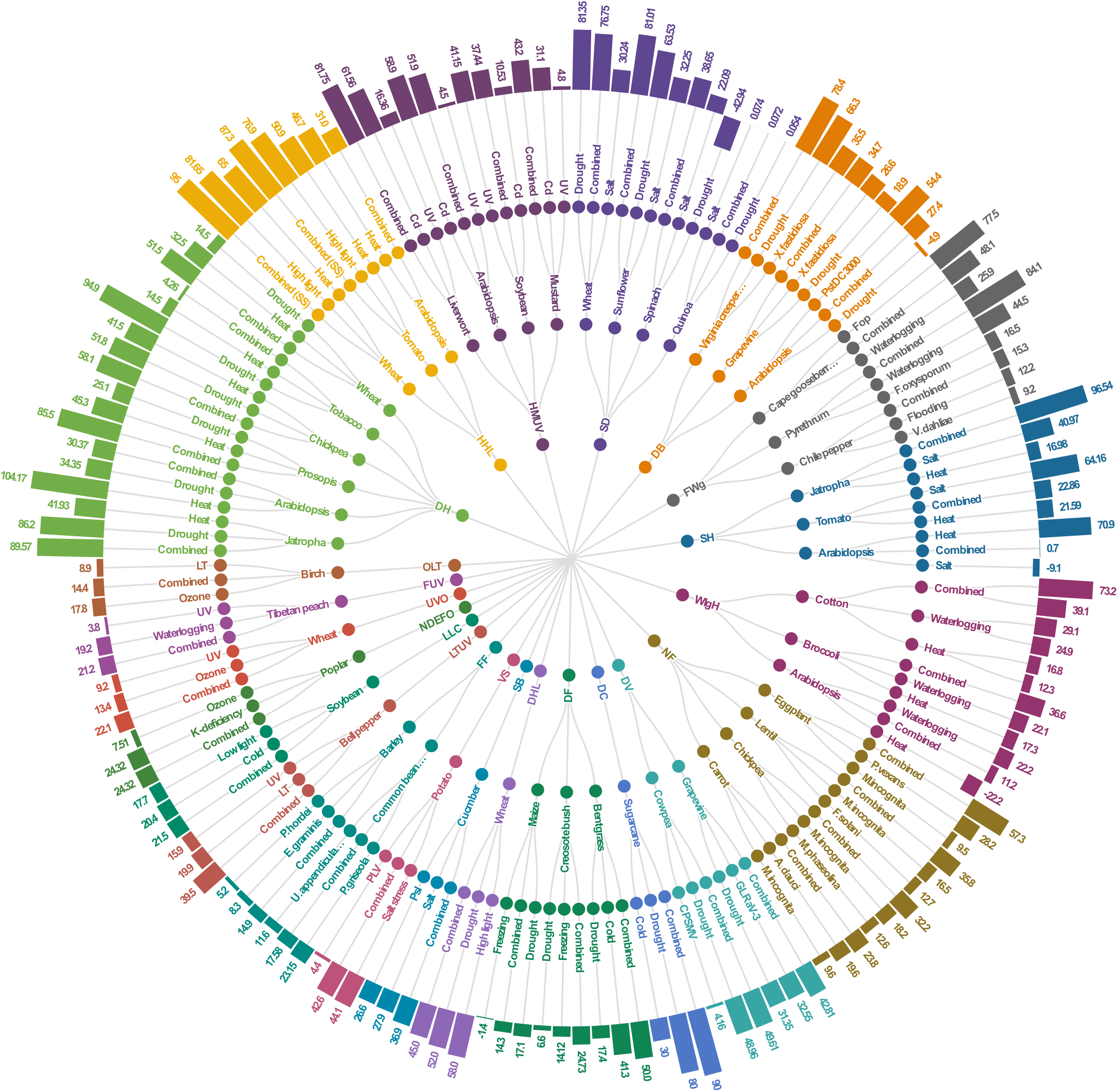
Literature analysis of combined stress physiological data on various plant species. The radial tree shows the effect of stress combination on physiological traits on various plant species. The tree was developed using Flourish studio (https://flourish.studio) and Tidyverse package in R (https://www.tidyverse.org/packages/). Traits included are photosynthesis, stomatal conductance, photochemical efficiency, Fv/Fm, and chlorophyll content. Percent under stress over control was calculated and using those values tree was developed. An interactive view of this tree is given on the SCIPDb website.

**Supplemental Figure 12.**
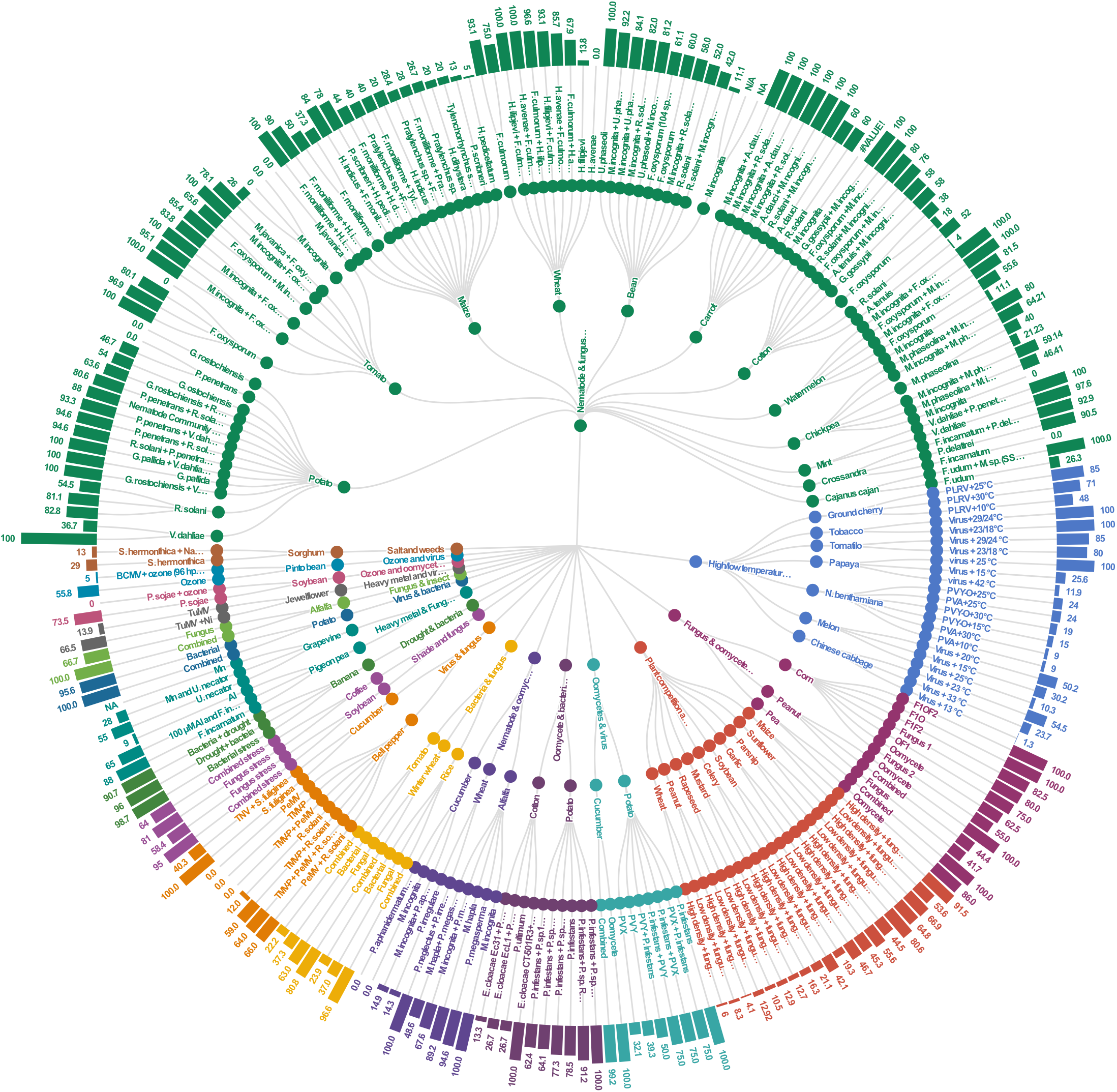
Literature analysis of disease incidence data under combined stress on various plant species. The radial tree shows the percent disease incidence in individual and combined stress conditions on various plant species. The tree was developed using Flourish studio (https://flourish.studio) and Tidyverse package in R(https://www.tidyverse.org/packages/). Organisms like bacteria, viruses, nematodes, fungus, mites, oomycetes, and insects were included in this analysis. Percent change in disease incidence under combined stress over individual stress was calculated and using those values tree was developed. An interactive view of this tree is given on the SCIPDb website.

## REFERENCES

Ahuja, I, de Vos, R.C., Bones, A.M., and Hall, R.D. (2010). Plant molecular stress responses face climate change. Trends in Plant Sci. 15:664–674.

Atkinson, N.J., and Urwin, P.E. (2012). The interaction of plant biotic and abiotic stresses: from genes to the field. J. Exp. Bot. 63:3523–3543.

Atkinson, N.J., Lilley, C.J., and Urwin, P.E. (2013). Identification of genes involved in the response of Arabidopsis to simultaneous biotic and abiotic stresses. Plant Physiol. 162(4):2028–2041.

Bjornson, M., Balcke, G.U., Xiao, Y., de Souza, A., Wang, J.Z., Zhabinskaya, D., Tagkopoulos, I., Tissier, A. and Dehesh, K. (2017). Integrated omics analyses of retrograde signaling mutant delineate interrelated stress-response strata. Plant J. 91(1):70–84.

Bian, Z., Gao, H. and Wang, C. (2020). NAC transcription factors as positive or negative regulators during ongoing battle between pathogens and our food crops. Int. J. Mol. Sci. 22(1):81.

Borkotoky, S., Saravanan, V., Jaiswal, A., Das, B., Selvaraj, S., Murali, A., and Lakshmi, P.T.V. (2013). The Arabidopsis stress responsive gene database. Int. J. Plant Genome 2013.

Cao, Y., Li, K., Li, Y., Zhao, X. and Wang, L. (2020). MYB transcription factors as regulators of secondary metabolism in plants. Biology 9(3):61.

Choudhary, A. and Senthil-Kumar, M. (2022). Drought attenuates plant defence against bacterial pathogens by suppressing the expression of CBP60g/SARD1 during combined stress. Plant Cell Environ. 45(4):1127–1145.

Cohen, S.P., and Leach, J.E. (2020). High temperature-induced plant disease susceptibility: more than the sum of its parts. Cur. Opin. Plant Biol. 56:235–241.

Cohen, I., Zandalinas, S.I., Huck, C., Fritschi, F.B., and Mittler, R. (2021). Meta-analysis of drought and heat stress combination impact on crop yield and yield components. Physiol. Plantarum 171(1):66–76.

Crandall, S.G., Gold, K.M., Jiménez-Gasco, M.D.M., Filgueiras, C.C. and Willett, D.S. (2020). A multi-omics approach to solving problems in plant disease ecology. Plos One 15(9):e0237975.

Desaint, H., Aoun, N., Deslandes, L., Vailleau, F., Roux, F., and Berthomé, R. (2021). Fight hard or die trying: when plants face pathogens under heat stress. New Phytol. 229(2):712–734.

Gupta, A., Sarkar, A.K., and Senthil-Kumar, M. (2016). Global transcriptional analysis reveals unique and shared responses in Arabidopsis thaliana exposed to combined drought and pathogen stress. Front. Plant Sci. 7:686.

Hamann, E., Blevins, C., Franks, S.J., Jameel, M.I., and Anderson, J.T. (2020). Climate change alters plant–herbivore interactions. New Phytol. 229:1894–1910.

IPCC Sixth Assessment Report: Climate Change 2022. https://www.unep.org/resources/report/ipcc-sixth-assessment-report-climate-change-2022

Li, X., Lawas, L.M., Malo, R., Glaubitz, U., Erban, A., Mauleon, R., Heuer, S., Zuther, E., Kopka, J., Hincha, D.K. and Jagadish, K.S. (2015). Metabolic and transcriptomic signatures of rice floral organs reveal sugar starvation as a factor in reproductive failure under heat and drought stress. Plant Cell Environ. 38(10):2171–2192.

Lopez-Delacalle, M., Silva, C.J., Mestre, T.C., Martinez, V., Blanco-Ulate, B., and Rivero, R.M. (2021). Synchronization of proline, ascorbate and oxidative stress pathways under the combination of salinity and heat in tomato plants. Env. Exp. Bot. 183:104351.

Lopez-Hidalgo, C., Guerrero-Sánchez, V.M., Gómez-Gálvez, I., Sánchez-Lucas, R., Castillejo-Sánchez, M.A., Maldonado-Alconada, A.M., Valledor, L. and Jorrín-Novo, J.V. (2018). A multi-omics analysis pipeline for the metabolic pathway reconstruction in the orphan species Quercus ilex. Front. Plant Sci. 9:935.

Mahalingam, R., Pandey, P., and Senthil-Kumar, M. (2018). Progress and prospects of concurrent or combined stress studies in plants. Ann. Plant Rev. 15:813–868.

Mittler, R. (2006). Abiotic stress, the field environment and stress combination. Trends Plant Sci. 11:15–19.

Mittler, R., and Blumwald, E. (2010). Genetic engineering for modern agriculture: challenges and perspectives. Annu. Rev. Plant Biol. 61:443–462.

Naika, M., Shameer, K., Mathew, O.K., Gowda, R., and Sowdhamini, R. (2013). STIFDB2: an updated version of plant stress-responsive transcription factor database with additional stress signals, stress-responsive transcription factor binding sites and stress-responsive genes in Arabidopsis and rice. Plant Cell Physiol. 54(2):e8–e8.

Pandey, P., Ramegowda, V., and Senthil-Kumar, M. (2015). Shared and unique responses of plants to multiple individual stresses and stress combinations: physiological and molecular mechanisms. Front Plant Sci. 6:723.

Pandey, P., Irulappan, V., Bagavathiannan, M.V., and Senthil-Kumar, M. (2017). Impact of combined abiotic and biotic stresses on plant growth and avenues for crop improvement by exploiting physio-morphological traits. Front Plant Sci. 8:537.

Pan, Q., Wei, J., Guo, F., Huang, S., Gong, Y., Liu, H., Liu, J. and Li, L. (2019). Trait ontology analysis based on association mapping studies bridges the gap between crop genomics and Phenomics. BMC Genomics 20(1):1–13.

Rasmussen, S., Barah, P., Suarez-Rodriguez, M.C., Bressendorff, S., Friis, P., Costantino, P., Bones, A.M., Nielsen, H.B. and Mundy, J. (2013). Transcriptome responses to combinations of stresses in Arabidopsis. Plant Physiol. 161(4):1783–1794.

Savary, S., and Willocquet, L. (2020). Modeling the impact of crop diseases on global food security. Ann. Rev. Phytopathol. 58:313–341.

Sinha, R., Irulappan, V., Patil, B.S., Reddy, P.C.O., Ramegowda, V., Mohan-Raju, B., Rangappa, K., Singh, H.K., Bhartiya, S. and Senthil-Kumar, M. (2021). Low soil moisture predisposes field-grown chickpea plants to dry root rot disease: evidence from simulation modeling and correlation analysis. Sci. Rep. 11(1):1–12.

Smita, S., Lenka, S.K., Katiyar, A., Jaiswal, P., Preece, J., and Bansal, K.C. (2011). QlicRice: a web interface for abiotic stress responsive QTL and loci interaction channels in rice. Database 2011.

Suzuki, N., Rivero, R.M., Shulaev, V., Blumwald, E., and Mittler, R. (2014). Abiotic and biotic stress combinations. New Phytol. 203:32–43.

Vemanna, R.S., Bakade, R., Bharti, P., Kumar, M.K., Sreeman, S.M., Senthil-Kumar, M. and Makarla, U. (2019). Cross-talk signaling in rice during combined drought and bacterial blight stress. Front. Plant Sci. 10:193.

Zandalinas, S.I., Balfagón, D., Gómez-Cadenas, A., and Mittler, R. (2022). Responses of plants to climate change: Metabolic changes during abiotic stress combination in plants. J. Exp. Bot. erac073, https://doi.org/10.1093/jxb/erac073.

Zandalinas, S.I., Fritschi, F.B., and Mittler, R. (2020a). Signal transduction networks during stress combination. J. Exp. Bot. 71(5):1734–1741.

Zandalinas, S.I., Fichman, Y., Devireddy, A.R., Sengupta, S., Azad, R.K., and Mittler, R. (2020b). Systemic signaling during abiotic stress combination in plants. Proc. Natl. Acad. Sci. USA 117(24):13810–13820.

Zandalinas, S.I., Fritschi, F.B., and Mittler, R. (2021a). Global warming, climate change, and environmental pollution: Recipe for a multifactorial stress combination disaster. Trends Plant Sci. 26(6): 588–599.

Zandalinas, S.I., and Mittler, R. (2022). Plant responses to multifactorial stress combination. New Phytologist. DOI: 10.1111/nph.18087.

Zandalinas, S.I., Sengupta, S., Fritschi, F.B., Azad, R.K., Nechushtai, R., and Mittler, R. (2021b). The impact of multifactorial stress combination on plant growth and survival. New Phytol. 230(3):1034–1048.

Zhang, H., and Sonnewald, U. (2017). Differences and commonalities of plant responses to single and combined stresses. Plant J. 90:839–855.

